# Towards comprehensive allosteric control over protein activity

**DOI:** 10.1101/384198

**Authors:** Enrico Guarnera, Igor N. Berezovsky

## Abstract

On the basis of the perturbation nature of allosteric communication, a computational framework is proposed for estimating the energetics of signaling caused by the ligand binding and mutations. The perturbations are modelled as alterations of the strenght of interactions in the protein contact network in the binding sites and neighborhoods of mutated residues. The combination of protein harmonic modelling with effect of perturbations and the estimate of local partition functions allow one to evaluate the energetics of allosteric communication at single residue level. The potential allosteric effect of a protein residue position, modulation range, is given by the difference between responses to stabilizing and destabilizing mutations. We show a versatility of the approach on three case studies of proteins with different mechanisms of allosteric regulation, testing it on their known regulatory and functional sites. Allosteric Signaling Maps (ASMs) obtained on the basis of residue-by-residue scanning are proposed as a comprehensive tool to explore a relationship between mutations allosterically modulating protein activity and those that mainly affect protein stability. Analysis of ASMs shows distance dependence of the mode switching in allosteric signaling, emphasizing the role of domains/subunits in protein allosteric communication as elements of a percolative system. Finally, ASMs can be used to complement and tune already existing signaling and to design new elements of allosteric regulation.

**Significance:** Universality of allosteric signaling in proteins, molecular machines, and receptors and great advantages of prospected allosteric drugs in highly specific, non-competitive, and modulatory nature of their actions call for deeper theoretical understanding of allosteric communication. In the energy landscape paradigm underliying the molecular mechanisms of protein function, allosteric signalling is the result of any perturbation, such as ligand binding, mutations, intermolecular interactions etc. We present a computational model, allowing to tackle the problem of modulating the energetics of protein allosteric communication. Using this method, Allosteric Signaling Maps (ASMs) are proposed as an approach to exhaustively describe allosteric signaling in the protein, making it possible to take protein activity under allosteric control.

## Introduction

Starting from phenomenological Monod-Weynman-Changeux (1) and Koshland-Nemethy-Filmer (2) considerations, the notion of allostery, i.e. modulation of protein activity by effector binding to remote regulatory exosites (3), had evolved into a well-formulated quantifiable concept (4). It is based on the key role of protein dynamics (5, 6) in providing the allosteric signaling regardless of the presence (7, 8) or absence (9, 10) of conformational changes in proteins of all structural types and functions (4, 11, 12). Non-competitive and modulatory nature of allosteric regulation together with high specificity of its signaling offers a great advantage for emerging type of medicines - allosteric drugs (13) – preventing toxicity effects, receptor desensitization (14), and providing functional selectivity (6, 7, 15). Design of allosteric drugs with desired agonist/antagonist activity requires in-depth understanding of allosteric mechanisms and ways of their adjustment (6, 7), resulting in rapidly growing number of experimental and computational studies (13, 16, 17).

The unifying theme of all kinds of observed allosteric regulation comes from its physical origin – perturbation of the protein dynamics. Ligand binding, mutations, post-translational modifications, intermolecular interactions, and their combinations can serve as a source of perturbation. Additionally, according to Cooper’s estimate that “*volume fluctuations of this order (30 cm^3^ mol^−1^) correspond to very small changes in overall dimensions of a globular protein, but if concentrated in one area would produce cavities or channels in the proteins sufficient to allow entry of solvent or probe molecules*”, allosteric communication can be even driven by pure equilibrium fluctuations (9). So far, perturbation-based approaches were directly used in the case studies of several proteins (18, 19). The whole-protein alanine-scanning mutagenesis showed that single mutation can also serve as a perturbation initiating and/or alternating allosteric signal (20), for example removal of critical residues disrupts the signal propagation in CFTR (21). Recent theoretical models and computational approaches describing the propagation of allosteric signal from a perturbation caused by the mutation/site are discussed elsewhere (22, 23).

As generic perturbation nature of allosteric signaling calls for developing common theoretical framework for quantitative analysis of allosteric signaling, we have recently proposed the structure-based statistical mechanical model of allostery (SBSMMA) that accounts for causality and energetics of allosteric communication upon ligand binding (24). In the SBSMMA, which is based on the harmonic modeling of a protein, the perturbation caused by the ligand binding is mimicked by increasing the stiffness of contacts between residues of the binding site. The model consists of three steps: (i) ligand free and ligand bound protein states are considered in the context of harmonic approximation and two sets of characteristic normal modes are obtained and used (ii) as an input for an allosteric potential, which evaluates the mean elastic work on a residue produced by its neighbors; (iii) partition functions characterizing the ensemble of all possible configurations of a residue neighbors are obtained and used for estimating the free energy difference between the ligand free and bound states. The latter is the configurational work exerted on a residue as a result of the change in the ensemble of configurations of its local neighbors induced through the perturbation. The model was also extended on considering the allosteric effect of mutations by modeling destabilizing/stabilizing mutations via weakening/strengthening the couplings in the contact network of a mutated residue (25, 26).

The goal of this work is to explore a possibility of obtaining comprehensive allosteric control over protein activity. To this end, we introduce here the notion of allosteric effect of mutation, which describes modulation of protein activity as a result of remote mutation. In order to have a generic description, we quantify the effect of mutation as a result of the change from the smallest (Ala/Gly-like) residue to the bulkiest one, regardless of the native amino acid in corresponding position. We start from the analysis of allosteric communication between known regulatory and functional sites, followed by the study of allosteric effect of mutations, combined impact of sites and mutations, and we finish with the comprehensive residue-by-residue scanning of allosteric effects of mutations and obtaining the Allosteric Signaling Maps (ASMs). ASMs allow to establish a clear relationship between mutations allosterically modulating protein activity and those that mainly affect stability. The change of mode in allosteric signaling is also observed in the analysis of ASMs as a function of distances, making it possible to discuss the role of domains/subunits in allosteric communication within multidomain/oligometic proteins and to pinpoint critical signaling inside and outside these structural units with the sign of modulation opposite to the one characteristic to these structural units. Using case studies of three proteins, Phosphofructokinase (PFK), D-3-Phosphoglycerate dehydrogenase (PGDH), and Insulin Degrading Enzyme (IDE), we investigate here the energetics of allosteric signaling caused by ligand binding, its cooperativity, and possibility to modulate it via allosterically acting mutations. We also show that ASMs provide an exhaustive information on allosteric communication in the protein, allowing, thus, to control protein activity in per-residue resolution by complementing and tuning already existing allosteric signaling with effects of mutations, as well as by designing new elements of regulation on the basis of allosteric signals of individual mutations and their combinations.

## Results

We propose here a theoretical model of allosteric control of protein activity aimed at quantification of the allosteric effects caused by ligand binding and mutations. Within the framework of the SBSMMA (24–26) the effect of a perturbation *P* (ligand binding, mutations, and their combinations) is evaluated at the single residue level as a free energy difference 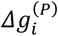 between any perturbed versus unperturbed protein states. The free energy difference 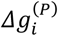 evaluates the change of the work exerted on the residue *i* as a result of the applied perturbation (see Equation 6 in Methods).

The portion of work exerted due to the purely allosteric effect is defined as the allosteric modulation 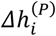, which, unlike 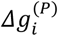, distinguishes between local and global effects that are induced upon a perturbation *P* (see Equation 8 in Methods). In other words, the allosteric modulation 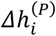 is considered as a background-free effect, where protein-average per-residue allosteric free energy is subtracted from the one detected on the residue of interest, showing the extent at which the allosteric signal to a corresponding residue/site is stronger than the background signaling between parts of the protein. The positive allosteric modulation is expressed in the increase of work exerted on a residue or regulated site, which may result in a local conformational change. The negative modulation, on the contrary, corresponds to a decrease of work, which may prevent conformational change.

In order to obtain a comprehensive allosteric control and conformational changes in a residue or regulated site and to be able to tune it (27), the allosteric effects caused by ligand binding are complemented with the substitutions of individual residues as basic elements of modulation. Fig. 1 shows a scheme of comprehensive allosteric control of protein activity, illustrating possible modes of allosteric regulation originated by different types of perturbations such as ligand binding (B, *Δh*^(*B*)^), stabilizing (UP, *Δh*^(*m*↑)^ and destabilizing (DOWN, *Δh*(*m*↓) mutations, and combinations of the ligand binding and mutations (*Δh*^(*Bm*↑)^ and *Δh*^(*B,m*↓)^. To consider a generic characteristic of the allosteric effect of mutations regardless of the natural residue in a selected position, a range of allosteric modulations *Δh*^(*m*↓↑)^ is obtained as a difference between the modulations obtained upon stabilizing (UP, substitution into bulky and strongly interacting residue) and destabilizing (DOWN, substitution into Ala/Gly-like residue) mutations. In this work, we explore the mechanisms of allosteric signaling in proteins of different structures, oligomeric states, and functions, considering tetrameric Phosphofructokinase (PFK), ring-shaped tetrameric D-3-Phosphoglycerate dehydrogenase (PGDH), and monomeric four-domain Insulin Degrading Enzyme (IDE).

**Fig. 1.**
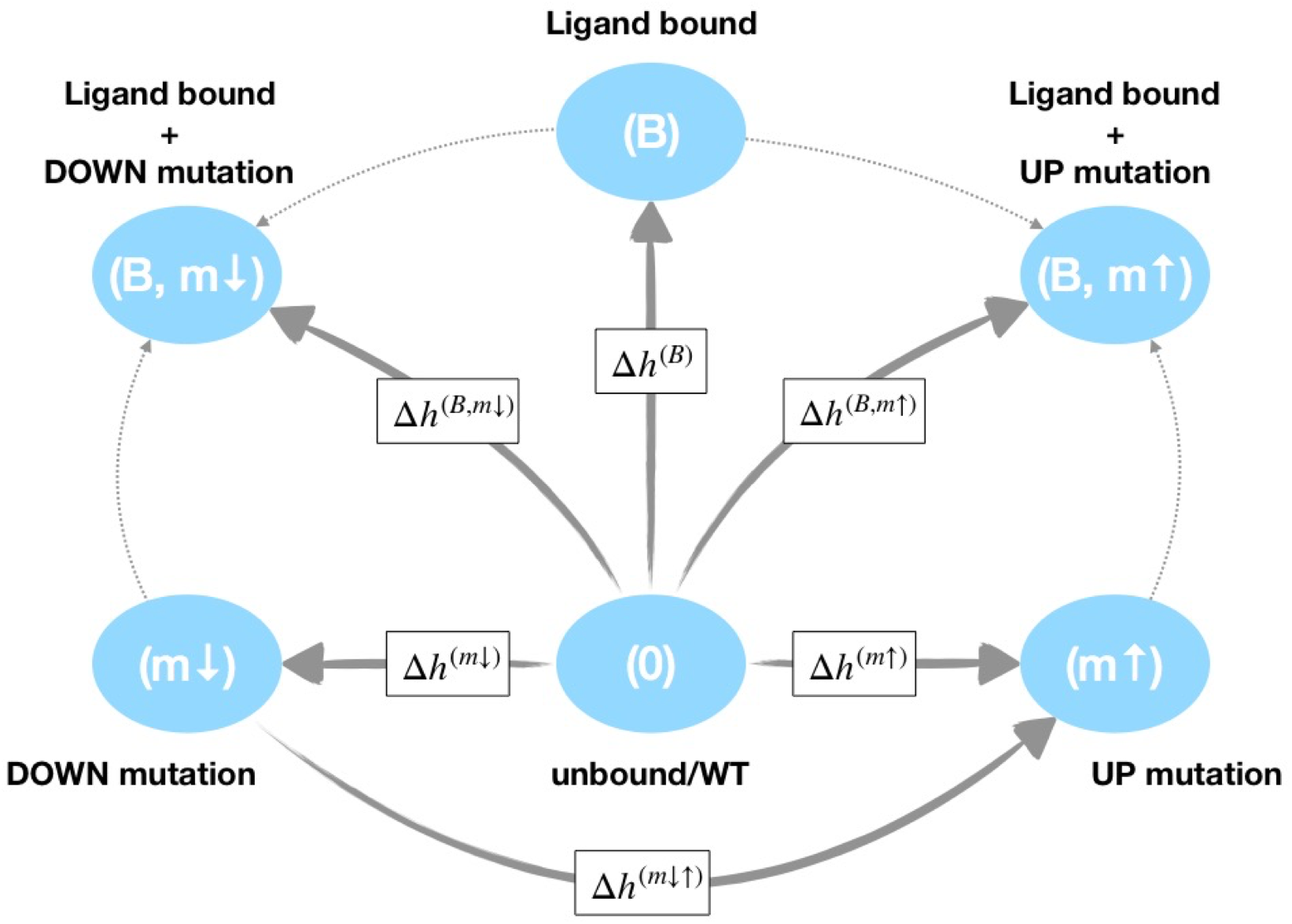
Comphrensive control of allosteric signaling. Allosteric modulations can be quantified for pairs of proteins states defined by different types of perturbations. Starting from a reference state (0, unbound/WT) the paired perturbed state is defined via either a ligand-bound state (B, *Δh*^(*B*)^, a mutated state (UP, *Δh*^(*m*↑)^ and DOWN, *Δh*^(*m*↓)^), or combined ligand-bound and mutated states (*Δh*^(*B,m*↑)^ and *Δh*^(*B,m*↓)^). The allosteric modulation associated with the transition from DOWN to UP mutated states identifies the allosteric *modulation range Δh*^(*m*↓↑)^, which is used as a generic quantity to evaluate the allosteric signaling upon mutation in the certain position in the protein regardless of the original amino acid in this position.

### Phosphofructokinase (PFK)

PFK phosphorylates fructose-6-phosphate (catalytic site F6P) in the process of glycolysis (28, 29), being activated by ADP (or GDP) and inhibited by phosphoenolpyruvate PEP. Fig. 2A shows a summarizing graph of the allosteric signaling in PFK originated by perturbation (simulated binding) in the corresponding binding sites of all four subunits of the tetramer (apo form was used, PDB ID: 3pfk). It shows that perturbation of the inhibitor (PEP) binding site induces a positive allosteric modulation at the catalytic site F6P 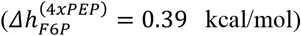 and a mild negative modulation at the substrate site ADPf 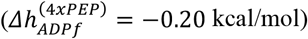. Similarly, perturbation at the activator (ADPa) binding site causes strong positive modulation at the catalytic site F6P 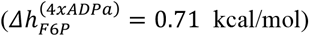 and a very weak negative modulation at the substrate site ADPf 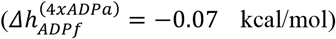. Binding sites of the activator ADPa and inhibitor PEP overlap significantly, and the difference in the result from their perturbation is expressed in the stronger positive modulation at the F6P functional sites and less negative modulation at the substrate binding site, ADPf, in the case of the activator (ADPa) binding. Therefore, one can assume that the mechanism of activation upon ADPa biding is based on the structural reorganization/adjustment resulting from the work exerted at the F6P site. PEP inhibitor, on the other hand, apparently acts by preventing the F6P and ADPf sites from achieving the functional state. Noteworthy, positive allosteric modulation at the functional 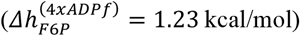 and both sites of effectors 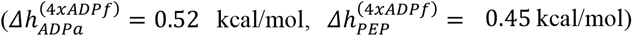 is also detected upon perturbation at the ADPf substrate site. To further explore this scenario of regulation, we consider the combined perturbation effect of ADPf substrate site together with either the ADPa-activator or PEP-inhibitor binding. In the case of activator, a stronger positive modulation on the catalytic site 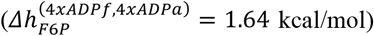 is detected rather than in the case when ADPf only is perturbed (+0.41 kcal/mol). When PEP and ADPf are bound together, modulation at the F6P site is unchanged compared to the effect of ADPf binding 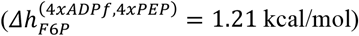. Altogether, one can conclude that ADPa acts with extra 0.41 kcal/mol in addition to 1.23 kcal/mol from ADPf, and the activating effect of ADPa binding is almost similar when it binds alone 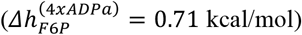 in comparison to the inhibiting PEP action 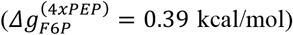.

**Fig. 2.**
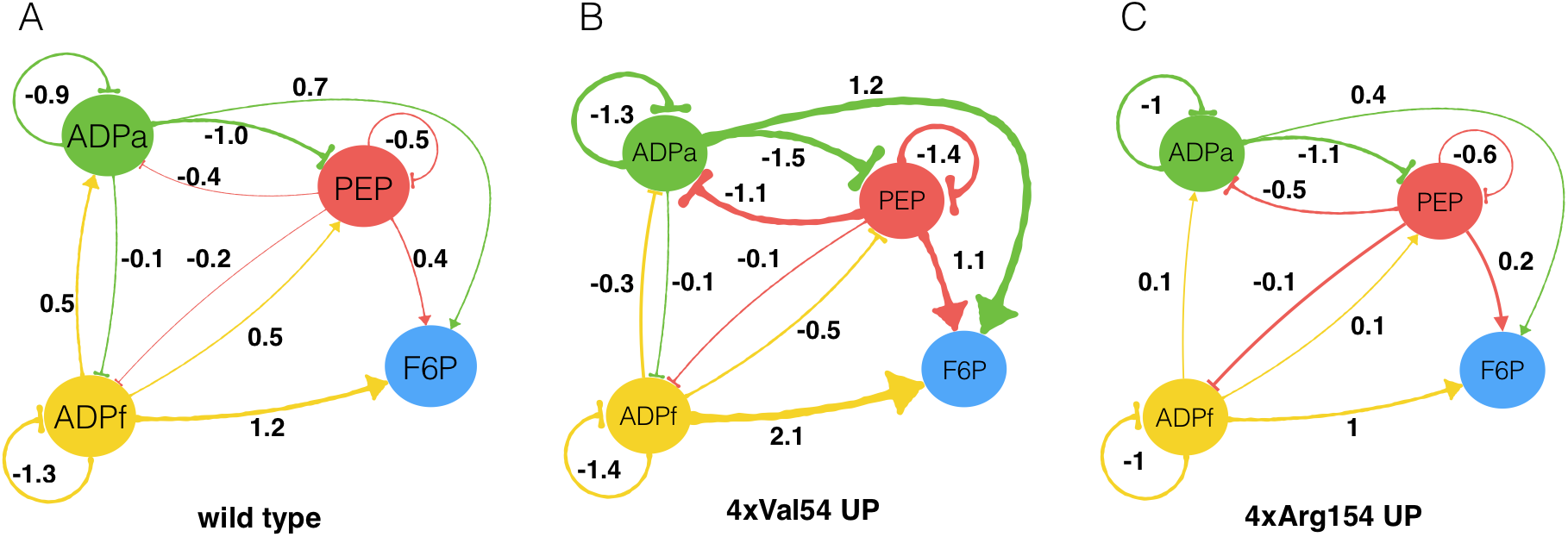
Allosteric communication in the phosphofructokinase (PFK). Graphs of allosteric signalling in the wild type (A) and mutated structures with UP mutations of Arg54 (B) and Arg154 (C). The apo-form of the PFK protein was used in calculations (PDB ID: 3pfk). Allosteric and functional sites are represented by nodes with linkers showing positive (arrows) and negative (bases) allosteric signaling. The thickness and the numbers on the arrows provide values of the allosteric modulation 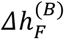 of the regulated site *F* upon the binding to regulatory exosite *B*.

The different nature of ADPa and PEP modes of action is also indicated in the cooperativity of F6P binding associated with the protein states with an increasing number of perturbed effector sites. Fig. S1A (see also Supplementary Table 1) shows the free energy response at the catalytic site 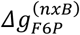 as a function of the number *n* of occupied effector sites (PEP or ADPa). The opposite concavities of the response curves suggest that PEP causes positive and ADPa – negative cooperativity in the F6P site. The mean cooperativity of the substrate binding (see Equation 7 in Methods) estimated for all pairs of protein states *n* and *n*+*1* upon sequential perturbation of PFK subunits is 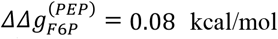 in case of PEP and 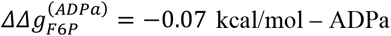.

Because only few residues make a difference between the binding sites of activator ADPa and inhibitor PEP, functional role of individual residues in these sites have been always a point of specific attention. Following earlier experimental work (30), we started from analyzing UP/DOWN mutations on the set of eight residues, Arg21, Arg25, Val54, Asp59, Arg154, Glu187, Arg211, and Lys213, belonging to the ADPa and PEP sites (Supplementary Table S2). Complete data on the effects of mutations on the functional and allosteric sites of PFK are presented in Fig. S2. For each residue, we considered protein states with single-residue mutations in one subunit and four-residue mutations in all subunits simultaneously, allowing to calculate corresponding modulation ranges *Δh*^(1 *x m*↓↑)^ and *Δh*^(4 *x m*↓↑)^. Mutations of residues Val54 and Arg154, yield strongest and, at the same time, opposite effect on the F6P site (Fig. 2B,C and Supplementary Table 2) upon mutations. Because of the difference in sizes of Arg and Val, modulation ranges obtained for these residues are 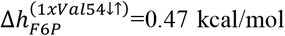 and 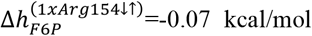 for single-residue and 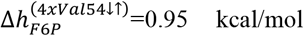 and 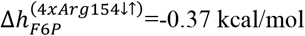 for four-residue mutations. Indeed, practically negligible negative modulation range of Arg154 can be explained by the bulkiness of native Arg154, which apparently allosterically prohibits the conformational changes in the functional F6P site. Mutation of Val54 shows, on the contrary, positive modulation range, pointing to the conformational changes in F6P site caused by the substitution into bulky amino acids.

Considering Val54 and Arg154 positions in PFK, we study a combined effect of allosteric modulation caused by the ligand binding and mutations. Strongly positive modulation range obtained for Val54 compared to weaker and negative one for Arg154 suggest to analyze the effect of stabilizing mutations of these residues on the allosteric signaling caused by the ligand binding (Fig. 1). Fig. 2B,C contains graph representations of the allosteric signaling associated with four-residue Val54 and Arg154 stabilizing mutations (UP), explaining details of the difference in the effects of these mutations in combination with ligand binding. Specifically, strong positive allosteric communication from all regulatory sites to F6P (*e.g*. 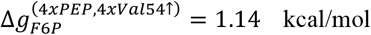 versus 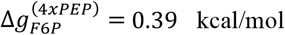 and 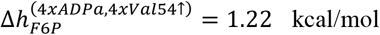 versus 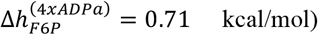), coupled with prevention of conformational changes that presumably repress binding to the regulatory ADPa and PEP sites as a result of ADPf perturbation (*e.g*. 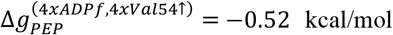 versus 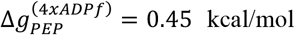 and 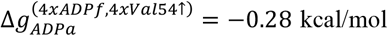 versus 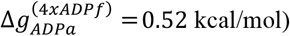), and no significant changes in signaling from the regulatory ADPa and PEP sites to ADPf (*e.g*. 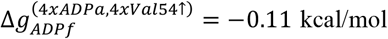 versus 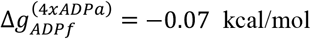 and 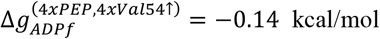 versus 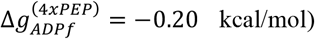). Thus, stabilizing mutation of Val54 residues shows that strong positive signaling to the functional site can be achieved via the recruitment of bulkier residues in place of Val54. Since UP mutation of Val54 strengthens positive signaling to F6P from both ADPa and PEP regulatory sites (Fig. 2B,C) along with positive modulation range 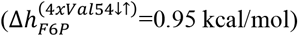 and wild-type positive modulation as a result of activator (ADPa) binding 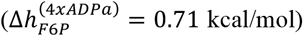, one can assume that it may change the mode of PEP site action from inhibiting to activating one.

A different picture of signaling in PFK is observed upon stabilizing mutations (UP) of the Arg154 residue, including some decrease in signaling from activating ADPa and substrate ADPf sites to F6P (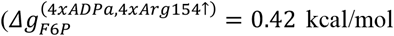 versus 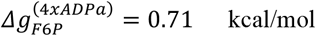 and 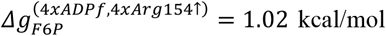 versus 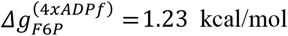, respectively), reduced signaling from PEP site to F6P (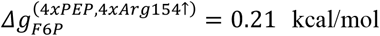 versus 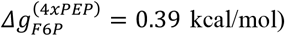), as well as from ADPf to PEP sites (Fig. 2). Since Arg154 is a bulky residue by itself, the only noticeable effect of its UP mutation is the decrease of communication between the PEP and functional F6P and ADPf sites, which may block an inhibiting signal from the site of the PEP inhibitor. At the same time, decreased positive allosteric modulation on F6P upon ADPa binding compared to wild-type regulation together with overall negative modulation simply show that bulky substitutions in Arg154, despite some repressive effect over the PEP site, are rather disruptive for overall allosteric signaling in PFK. Finally, opposite weak cooperativity, positive in the case of Val54 and negative in Arg154 mutations (Fig. S1B and Table S1), respectively, is observed, further supporting the difference in the ways of modulation of allosteric signaling in PFK caused by these mutations.

### D-3-Phosphoglycerate dehydrogenase (PGDH)

The ring-shaped tetrameric PGDH is an allosterically regulated enzyme that catalyzes formation of 3-phosphohydroxypyruvate from 3-phospho-D-glycerate with NAD as cofactor. Contrary to the majority of allosterically regulated enzymes PGDH is a V-type enzyme, that is the rate of reaction but not the substrate binding is allosterically modulated (31). Each PGDH subunit contains three well defined structural domains: the cofactor binding domain (NAD sites), the substrate binding domain (AKG sites), and the regulatory domain. PGDH is allosterically inhibited by L-serine (SER site) via binding at the interface between the regulatory domains of adjacent units. Structural analysis suggests that PGDH catalytic activity and its allosteric inhibition occurs through the hinge-mediated rigid motion of domains (31). The hinges H2 and H3, as the caps of helix Gln298-Asn318, and additional hinge H4, constitute a pin shaped spring, which performs the function of a signal transferring device from the regulatory to the substrate domains (Fig. 3A,B). The L-serine binding causes simultaneous opening of the interface between adjacent regulatory domains and of the active site cleft, inhibiting the catalytic activity (32). Our calculations (active form was used, PDB ID: 1yba) show that upon binding at the SER effector site the protein domains respond with different modes of allosteric modulation: it is positive in the effector 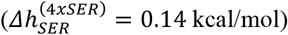 and cofactor 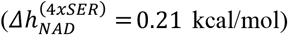 domains, but weakly negative in substrate-binding one (AKG, 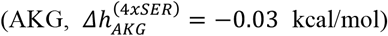). These results corroborate the modular response of PGDH discussed elsewhere (31), which is in agreement with the idea that SER binding induces a more open form of the interface between regulatory domains (33) reflected in a positive modulation observed in regulatory domains. Additionally, weak allosteric modulation detected at the substrate site AKG can apparently serve as an indication of the V-type nature of PGDH enzyme (31) with modulation of the reaction rate, but not the substrate binding being affected (32).

**Fig. 3.**
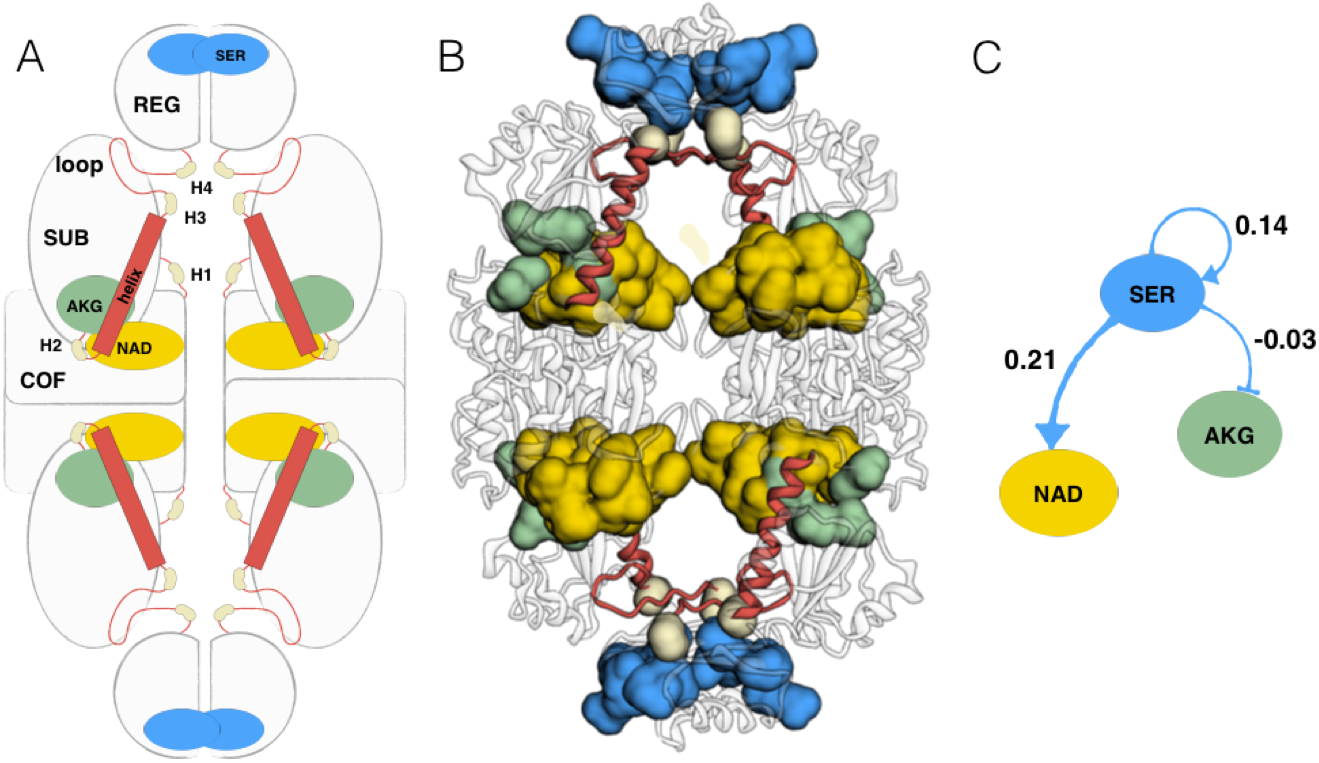
Domain-based allosteric regulation in D-3-phsophoglycerate dehydrogenase (PGDH). (A) Cartoon representation of the PGDH with indicated hinges and helices, which are crucial for allosteric signalling between regulatory (contains SER-binding site), substrate (AKG), and cofactor (NAD) domains. The hinges are located between the cofactor binding domain and the substrate binding domain (H1: Pro105-Phe106), and between the substrate binding domain and the regulatory domain (H2: Gly294-Gly295, H3: Gly319-Ser320, and H4: Gly336-Gly337). (B) Ribbon representation of the PGDH, showing allosterically-relevant structural connections (helices and hinges) between regultory and fucntional sites. (C) Graph representation of the allosteric signaling in PGDH upon the L-serine binding.

The major question we address in PGDH analysis, is whether and to what extent mutations can affect domain-based mode of allosteric communication and regulation of protein activity. Complete data on the effects of mutations on the functional and allosteric sites of PGDH is presented in Fig. S3. We specifically considered mutations in hinges, H1 (Pro105, Phe106), H2 (Gly294, Gly295), H3 (Gly319, Ser320), and H4 (Gly336, Gly337), and in the SER-binding site (Ile365 and Leu370), see Table 1. Mutations of residues in H1 result in the negative modulation range in the cofactor and substrate domains (Fig. 3), which suggest an overall decrease of motion of these domains around hinge H1. At the same time, mutations of Phe106 originate some structuring/orientation in the regulatory site caused by the positive modulation 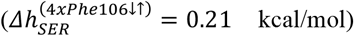. Mutations of the hinge H2 do not significantly affect neither of the sites/domains. Both hinges H3 and H4 are located between the regulatory and cofactor domains before the helix of pin-shaped spring, which is crucial for conformational changes involving these domains. Mutations of residues in these hinges lead to the substantial allosteric modulation of PGDH response caused by the L-serine binding (Fig. 3): (i) they result in positive modulation on the cofactor domain and negative modulation on the regulatory sites (very strong in case of mutations in H4); (ii) mutations in H4 change the sign of modulation on the substrate binding site compared to the effect of the L-serine binding, resulting in a positive work exerted on it in the case of mutations in all four subunits. All the stabilizing mutations (Gly319/H3, Gly336/H4, and Gly337/H4) apparently lead to canceling the inhibition mode, as they can prevent binding of the inhibitor (negative modulation, 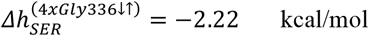 and 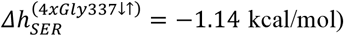) and, on the contrary, can enable binding of the substrate (positive modulation, 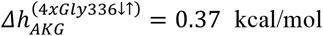 and 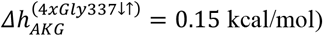). Additional positive work exerted in the cofactor site can additionally support binding of (NAD 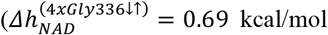 and 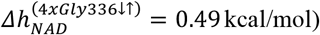). Noteworthy, three of the four mutated residues are glycines, hence the effect of mutation is expected to be more pronounced especially in cases of four-residue mutation (Ser320/H3 mutation shows the weakest effect, see Table 1). Finally, we compared effects of two mutations in the regulatory site, Ile365 and Leu370, which despite their close location to each other, size, and similar physical-chemical characteristics modulate allosteric signaling in the opposite way. Mutations of Ile365 induces a weak positive modulation in the substrate site 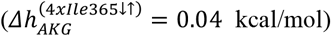 and strong negative - in the regulatory one 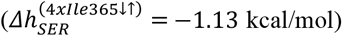, while just the opposite happens in case of Ile370 mutation (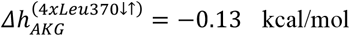 and 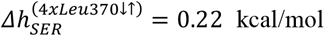). This, in addition to the importance of single-residue mutations in modulation of allosteric signaling observed for positions in difference hinges, a drastic difference in the results of mutations of similar and closely located Ile365 and Ile370 shows that any residue position in the protein can potentially be a source of allosteric modulation.

**Table 1:**
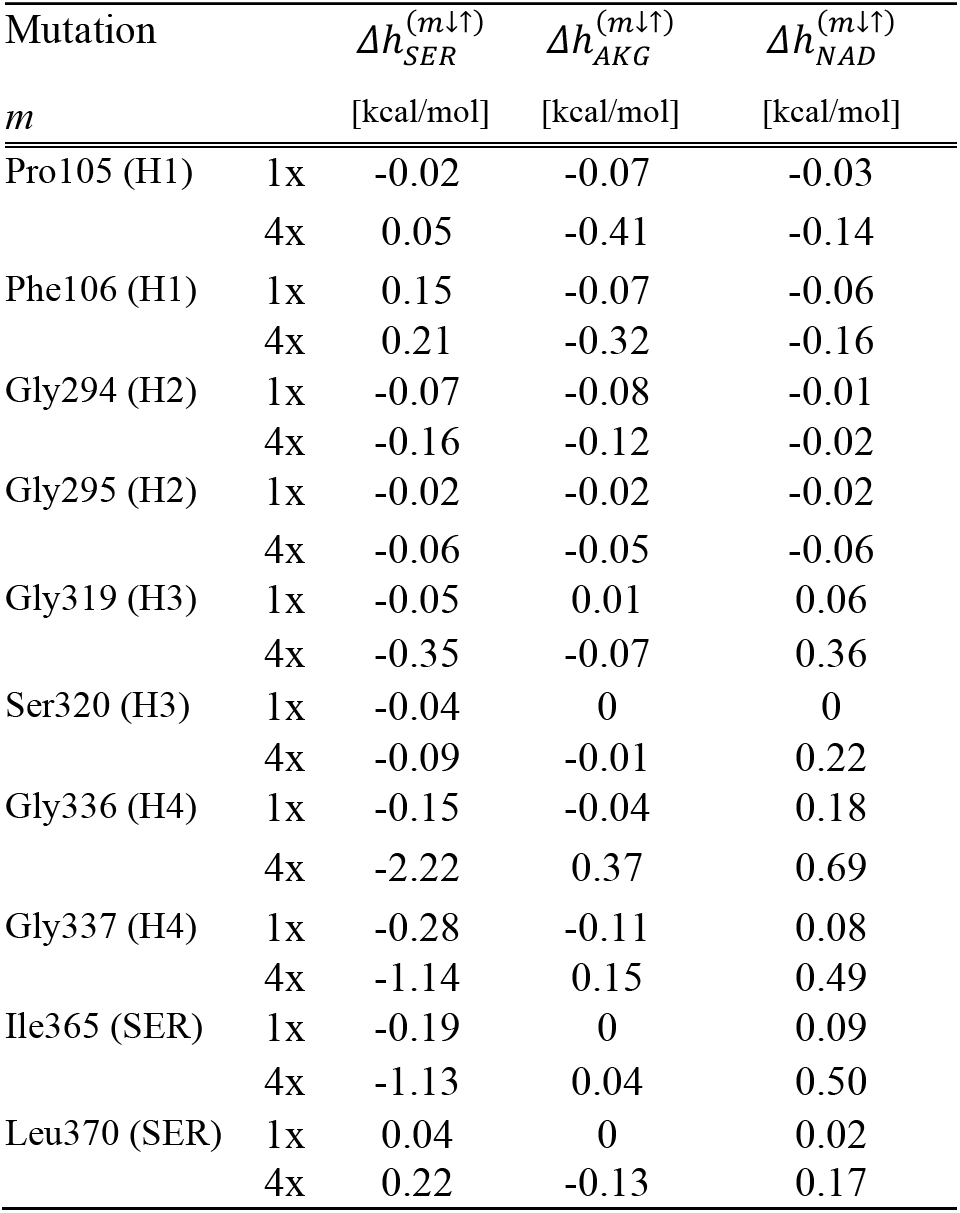
The list of residues mutated in PGDH protein. Single-residue (1x) and four-residue (4x) UP/DOWN mutations are considered and the corresponding difference of allosteric modulations between the UP and DOWN mutated states *Δh*^(m↑)^ = *Δh*^(*m*↓)^ − *Δh*^(*m*↓)^ - modulation ranges - are reported for the sites SER, AKG, and NAD.

### Insulin Degrading Enzyme (IDE)

Insulin Degrading Enzyme is a monomeric Zn^2+^-dependent protease organized in four domains, with the interface between the amino- (domains 1 and 2) and carboxy-terminal (domains 3 and 4) halves forming the degradation chamber. The cleavage (CLS), Zn^2+^-binding (ZN), and β-recognition sites (AB) are spatially close in the structure, constituting the complex active site where the substrate binding, recognition, and hydrolysis take place. The substrate acquisition is supported by its anchoring in the additional exosite (EXO) (34), and its proteolysis is allosterically regulated by the ATP bound to remote (ATP) site (35). Though insulin was the first molecule recognized as the substrate, hence Insulin-Degrading Enzyme (IDE), it appeared that this protein has a unique specificity towards β-structure forming aggregation-prone molecules, including Aβ and other amyloidogenic peptides, hormones, chemokines, and growth factors to name a few (36). It is of great importance for the context of this work that in addition to IDE’s allosteric effector ATP, protein activity can be also allosterically modulated and targeted against specific substrates by different molecular groups, small molecules, post-translational modifications, and mutations (25, 36).

Since the exosite (EXO) is critical for anchoring substrates and the ATP-binding site (ATP) provides allosteric modulation of IDE activity, we first explore effects of binding to these sites on the catalytic (CLS), Zn^2+^(ZN), and β-recognition (AB) sites (Zn-bound form of IDE in complex with insulin B chain was used, PDB ID: 2g54, see Fig. 4A). In general, perturbation of the EXO and ATP sites provides rather modest allosteric signaling to the catalytic sites (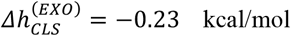 and 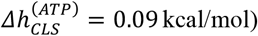), except the relatively high positive modulations of ZN (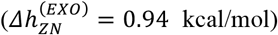 and AB 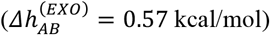) sites upon the EXO-site’s perturbation. Positive modulation on the AB and ZN sites becomes stronger 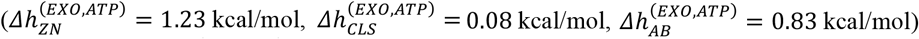 when both EXO and ATP sites are perturbed, pointing to the importance of mutual effect of binding to these sites for modulation of the catalytic site activity (see Fig. 4B). Binding to these sites also allosterically affects dynamics of domains, which was shown to be important for IDE activity and its modulation (25). Perturbation of the EXO site causes positive modulation in domains 1 and 4 (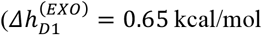 and 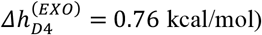) and negative modulation in domains 2 and 3 (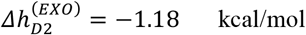 and 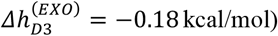). ATP binding originates a mild positive modulation in domain 2, apparently facilitating binding to the EXO site located in this domain (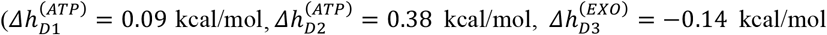, and 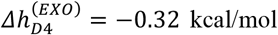). Mutual binding to EXO and ATP sites leaves positive modulation only on domain 1 (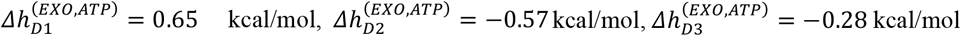, and 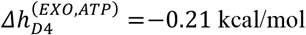), characterizing state of the protein in which the substrate is bound, and it is being processed.

**Fig. 4.**
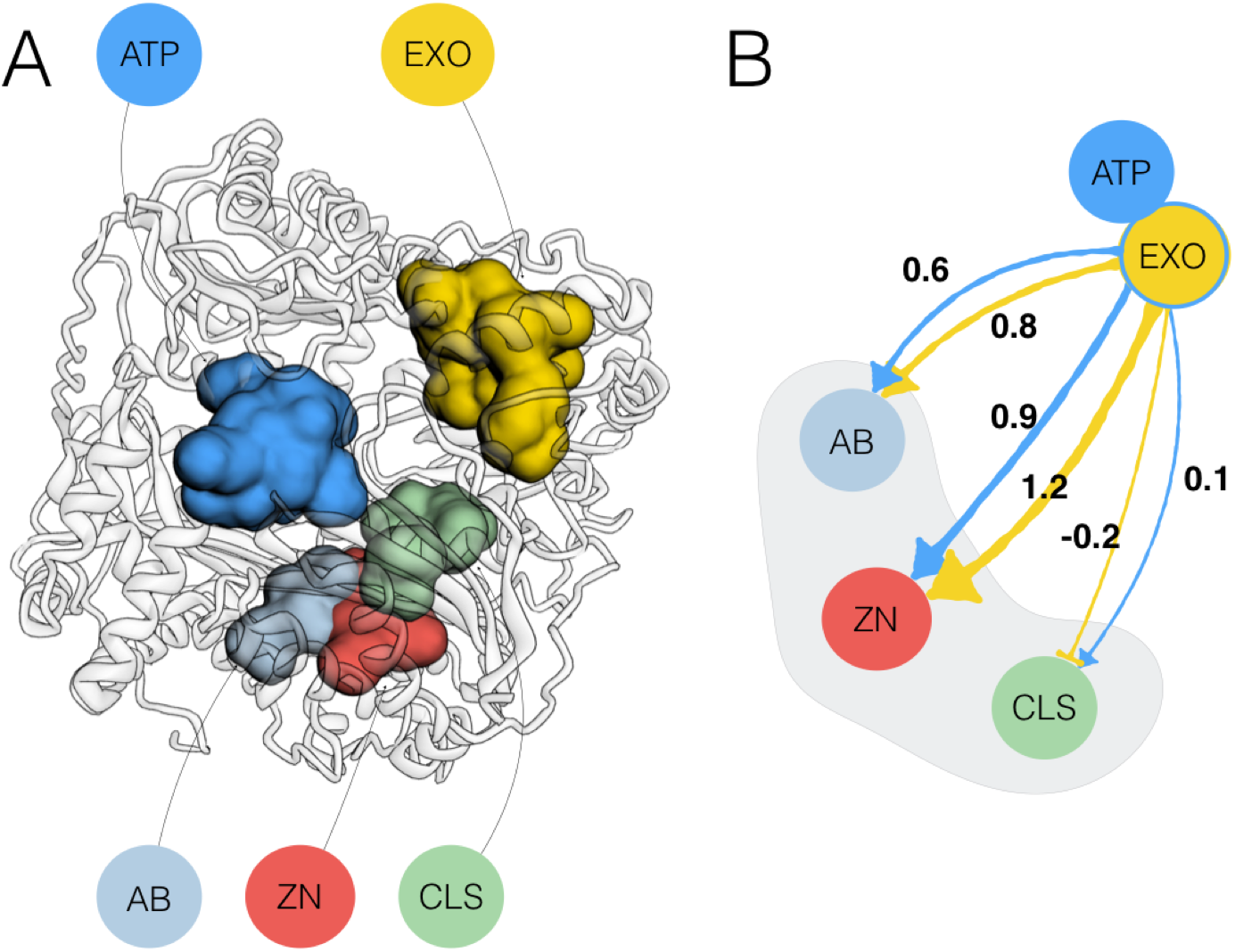
Allosteric signalling between functional and regulatory sites of Insulin-Degrading Enzyme (IDE). (A) Ribbon representation of IDE shows location of functional and regulatory sites: AB - recognition site specific to detecting Aβ peptide and other β-structure forming aggregation-prone substrates; CLS - substrate cleavage site; EXO - exosite anchoring the IDE’s substrates; ZN - Zn^2+^-binding site; ATP – ATP-effector binding site. (B) Allosteric signalling from ATP and EXO sites to the componenets of the functional site (AB, CLS, and ZN). The single-chain IDE consists of four domains: residues 43-285 (Domainl), 286-515 (D2), 542-768 (D3), and 769-1016 (D4). Functional and regulatory sites of IDE include: ZN (D1: His108, His112, Glu189); CLS (D1: Phe141, Trp199, Phe202); AB (D4: Arg824, Tyr831); EXO (D2: His332, His336, Leu337, Ile338, Gly339, 340, His341, Leu359, Val360, Gly361, Gly362, Gln363; D3: Tyr609); ATP (D2: Arg429; D4: Asp895, Lys896, Pro897, Lys898, Lys899, Ala902, Ala905).

Recent studies of five allosteric mutations, including both computational analysis and experimental verification, showed that specific modulation of IDE’s activity against Aβ substrate can be achieved by allosterically affecting the energetics of domains and components of the functional site (25). It seems promising, therefore, to use allosteric effects of mutations for activation of IDE against targets of interest, leaving its activity against other substrates unchanged. As the first step in this direction we obtained here a complete data on the allosteric effect of mutations on the energetics of functional and regulatory sites (sites: EXO, CLS, ZN, AB, ATP; Fig. S4) and on the modulation of domains (Domains 1-4; Fig. S5). The modulation range obtained for each position is the difference between the effect of the stabilizing and destabilizing mutations, which quantifies the allosteric modulation change upon the switch from the smallest to the bulkiest residue in a certain position regardless of the original native amino acid in this position. Therefore, within our model, mutations ranging from destabilizing to stabilizing can partially mimic the effect of binding in the environment of the corresponding residue. Mutations in positions located in ATP and EXO sites clearly show an activating nature of these sites on IDE activity (see the modulation range obtained from destabilizing to stabilizing mutations shown in Fig. S4). In particular, all elements of the functional site (AB, CLS, EXO, and ZN) yield positive allosteric modulation range – indicator of conformational changes facilitating protein activity as a result of allosteric signaling from the ATP site. Mutations in the exosite EXO provide positive modulation range in AB, ATP, and ZN sites, and negative in CLS, which apparently points to the importance of recognizing β-structure forming substrate before its processing in the CLS site (Fig. S4). Mutations in AB and ZN sites induce a positive modulation range in ATP and EXO sites. Because of their relative closeness (see Fig. 4A and Fig. S4 and S6C), these sites work against each other and the CLS site, which also suggests that the sequence of events in the substrate processing concludes with the involvement of CLS site. Sequential mutations in IDE domains show a consistent picture of positive modulation range in the diagonal domains, *i. e*. in the pairs of domains 1-3/3-1 and 2-4/4-2 (Fig. S5). Mixed positive-negative modulation range is systematically observed in domains flanking domain n, namely domains (n-1) and (n+1), in which mutations are performed. The patterns of mixed modulation range in flanking domains reflect a structural similarity between IDE domains, which are all homologous aβ roll folds. At the same time, all domains share at maximum 15% of sequence identity, which is reflected in numerous local diversity of observed modulation both in terms of the sign and the value.

### Exhaustive analysis of allosteric signaling: Allosteric Signaling Maps

The analysis of three case studies proteins, PFK, PGDH, and IDE, shows that despite structural and functional differences allosteric signaling is inherent in these proteins, it can be generically described using formalism of modulation in per residue approximation, and it can be quantified for functional and regulatory sites, (sub)domains, chains, and other structural units. Fig. 5 contains distributions of allosteric modulation ranges as a result of point mutations Val54 in PFK, Gly33 in PGDH, and Phe807 in IDE, respectively. Positive allosteric modulation is exemplified by the sites F6P, NAD, and EXO, as well as by individual residues (200 in PFK and PGDH, and 500 in IDE) in these proteins. Though, in general, distributions are skewed to the positive values, their shapes and ranges of negative/positive modulations are specific for each protein and depend on the structural characteristics, such as oligomerization state, types of folds/domains forming the protein, linkers/hinges between them. Mutations of small residues, such as Val54 (PFK) and Gly334 (PGDH) are expected to result in a stronger modulation imposed on responding sites and residues as well as stronger local negative modulation around mutated residues compared to substitution of rather bulky Phe807 (IDE). In other words, the sequence dependence of allosteric modulation is also at play in addition to structure dependence shown above. Therefore, to obtain the complete picture of regulation and to be able to consider any individual or combined effect(s) of the ligand(s) binding and/or mutation(s) in the context of structure-function relationship one has to perform an exhaustive scanning of mutations and obtain their sequence-structure dependent modulatory effects.

**Fig. 5.**
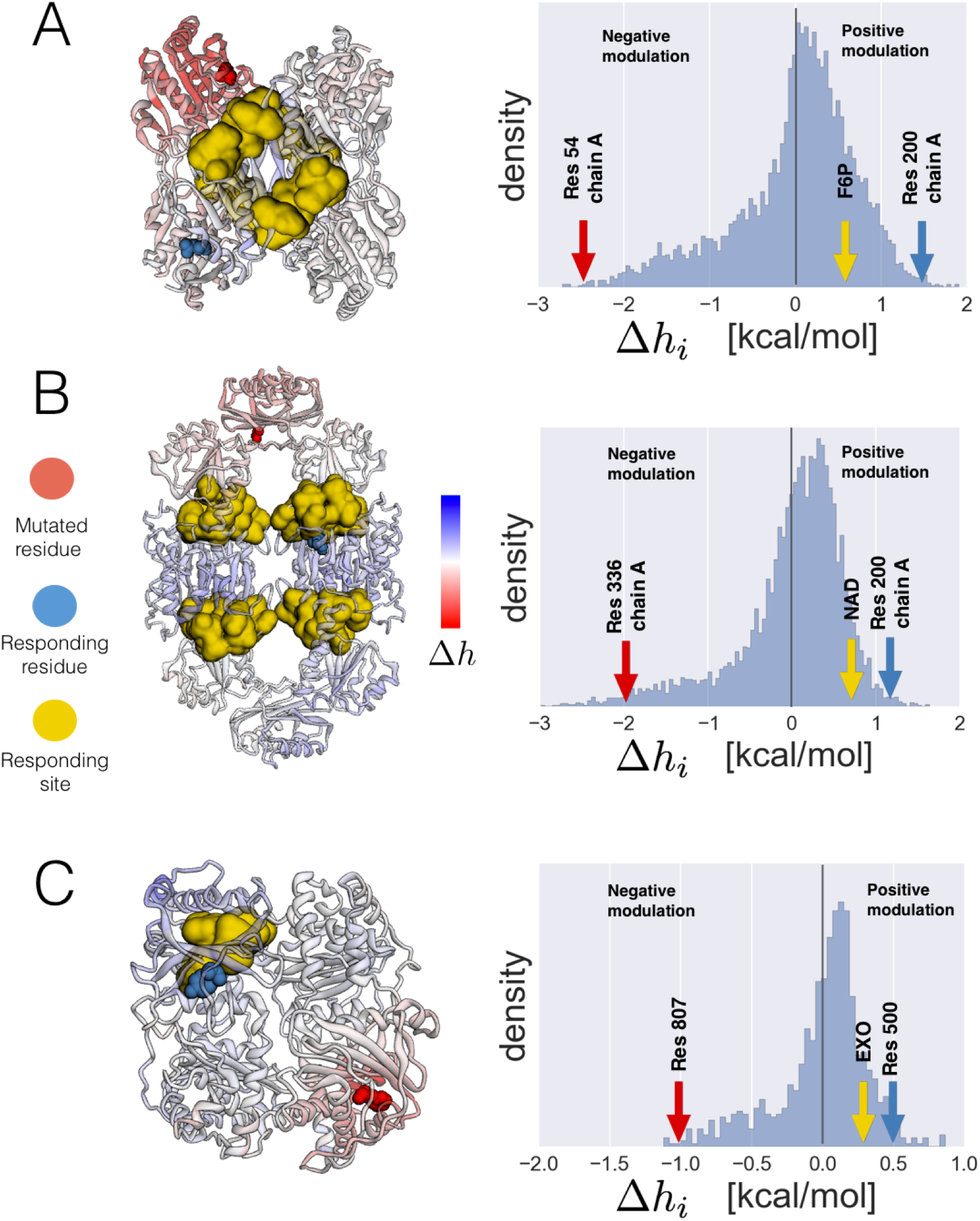
Distributions of allosteric modulation ranges caused by single-residue mutations in PFK, PGDH, and IDE. (A) Mutation of residue 54 in PFK; (B) Mutation of residue 336 in PGDH; (C) Mutation of residue 807 in IDE.

The calculation of the allosteric modulation range 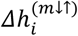 (Equation 11, Methods) upon residue-by-residue mutations provide the complete pattern of allosteric response upon perturbation, which we call Allosteric Signaling Map (ASM). Because of the nature of the allosteric potential, i.e. elastic energy applied to a residue depends on the configurational states of the neighboring residues, the ASM is an inherently asymmetric matrix. Fig. 6 contains ASMs for the proteins studied in this work, PFK (panel A), PGDH (B), and IDE (C), respectively. Major patterns of negative and positive modulations are clearly distinguishable in ASMs, revealing the domain and oligomeric structures of proteins. Specifically, negative modulation ranges delineate compact domains and monomers located along the diagonals of corresponding ASMs. In tetrameric PFK each monomer consists of two domains strongly interacting with each other via last quarter of the second domain (Fig. 5A). Each of the PGDH subunits consists of cofactor, substrate, and regulatory domains clearly visible along the matrix diagonal (Fig. 5B). Four domains of the single-chain IDE also form a characteristic pattern of negative modulation (Fig. 5C).

**Fig. 6.**
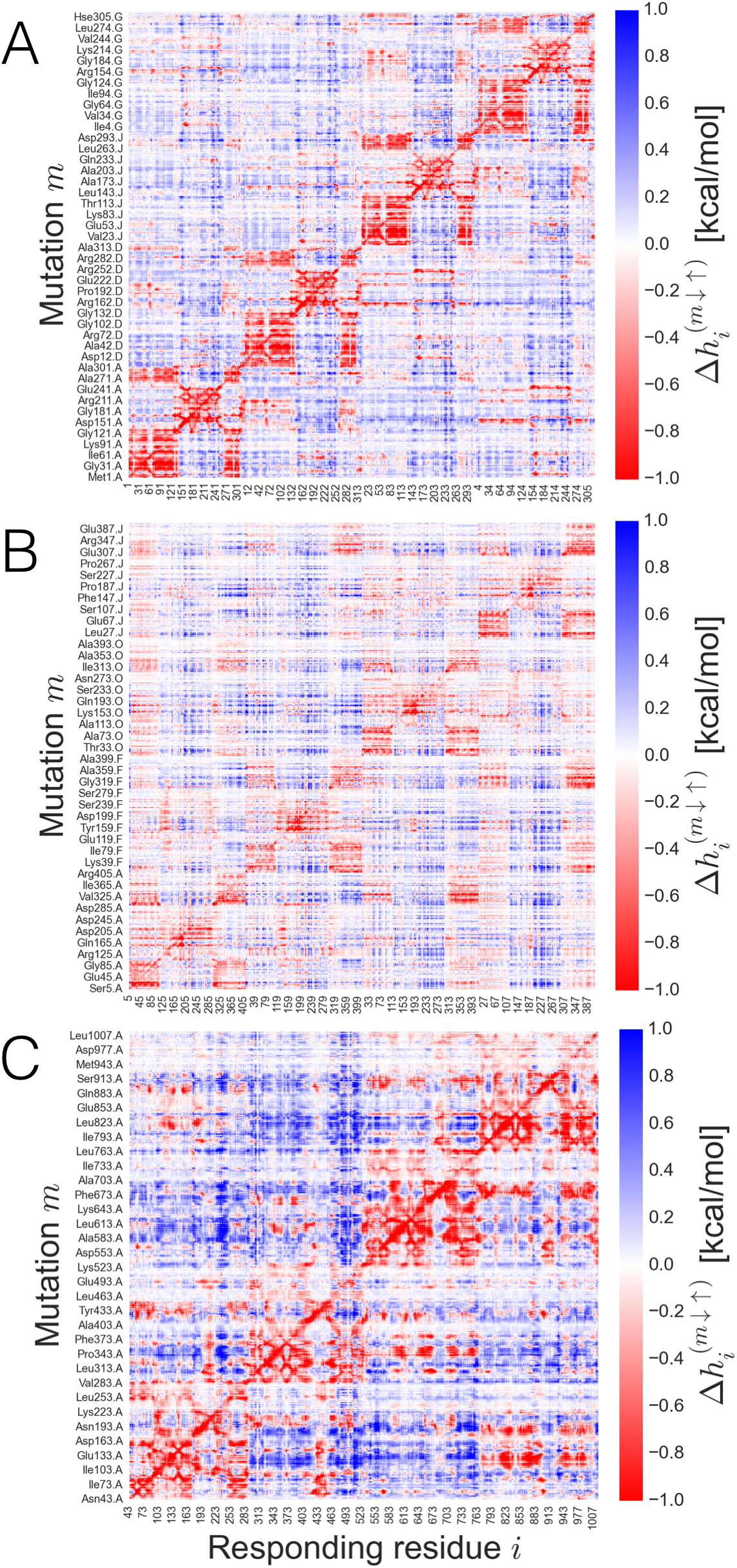
Allosteric Signaling Maps (ASMs) for the protein PFK (A), PGDH (B), and IDE (C). The allosteric modulation caused by the generic mutation *m* (y-axis) on a responding residue *i* (x-axis) is expressed via the modulation range 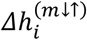. The modulation range is obtained as a difference between the modulations by the UP (stabilizing) 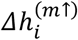 and DOWN (destabilizing) 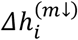 mutations, respectively. The modulation range 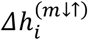 provides the energetics of the allosteric response caused by mutation/perturbation ranging between two extreme states. Negative modulations (red) indicate that upon generic mutation the responding residues experience a decrease of work exerted on them. On the contrary, positive modulations (blue) indicate that responding residues experience an increase of the work, leading to conformational changes.

Chiefly negative modulation range within compact structural units (distance matrices are shown in Fig. S6) is a consequence of different effects of UP and DOWN mutations (Fig. 6). UP mutations dominate the modulation, varying from strongly negative typical for over-stabilizing interactions of the mutated residue environment to weakly positive at longer distances (Fig. 6 and Fig. S6). The change of sign in modulation by UP mutations and almost complete decay from originally weak signal from DOWN mutations results in the change of the modulation mode from negative to positive at the distances 29 Å in PFK, 40 Å in PGDH, and 34 Å in IDE, respectively (Fig. 6 and Fig. S6). These critical lengths at which allosteric signaling changes its sign match to the typical radius of gyration for protein domain sizes ranging from 50 to 350 residues (37). Remarkably, recent analysis of chemical shift perturbation datasets originated by the ligands binding and mutations (38) revealed a universal distance-dependent decoy of the allosteric signal percolation within 20-25 Å. Fig. 6 and Fig. 7 supported by the above experimental data emphasize, therefore, on the two-faceted nature of allosteric modulation: (i) dominating negative modulation acts inside compact structural units where perturbation takes place, causing (ii) mostly positive modulation in other parts of the protein starting at the borders of perturbed monomers/domains, percolating through the rest of the protein as a result of the low frequency normal mode dynamics. It should be noted, however, that modulation of opposite signs observed in both cases points to the importance of corresponding signaling inside and outside of compact structural units.

**Fig. 7.**
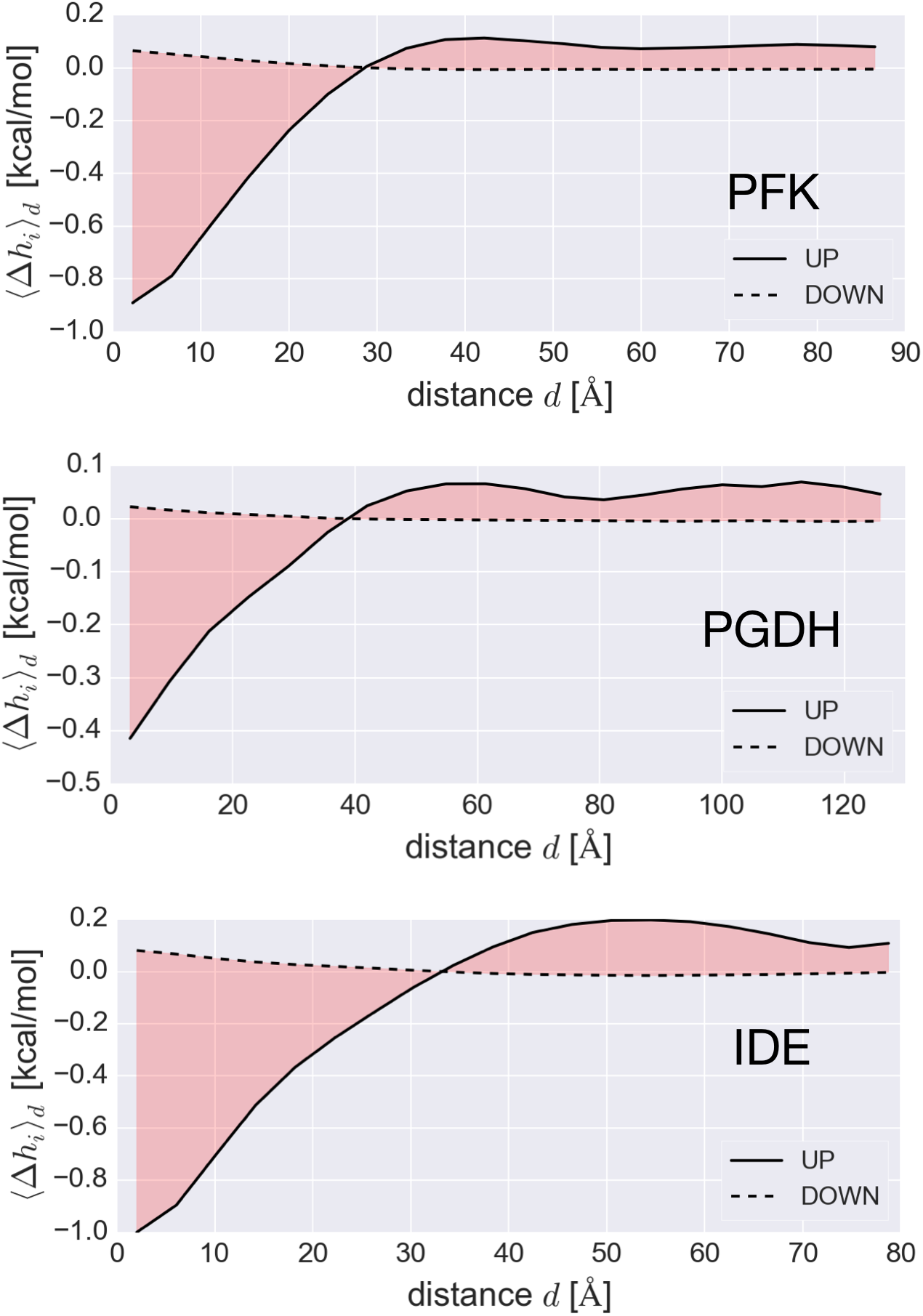
Distance-dependence of allosteric modulation in in PFK (A), PGDH (B), and IDE (C). The UP mutations cause strong negative modulation at short distances from the perturbation, starting from –0.93 kcal/mol in PFK, –0.42 kcal/mol in PGDH, and –0.98 kcal/mol in IDE, which converges towards positive residual value (0.11 kcal/mol in PFK, 0.06 kcal/mol in PGDH, and 0.09 kcal/mol in IDE). The modulation caused by DOWN mutations is generally weak and positive (starts from 0.06 kcal/mol in PFK, 0.02 kcal/mol in OGDH, and 0.08 kcal/mol in IDE), decaying to a very small residual value. A critical length for the propagation of allosteric signals can be identified from the distance where the mean allosteric modulation for UP and DOWN mutations switch their mode (29 Å in PFK, 40 Å in PGDH, and 34 Å in IDE).

The signaling observed between different sites of proteins (Fig. 2A, Fig. 3C and Fig. 4B), its modulation by mutations (Fig. 2B,C and Supplementary Table S1), and cooperativity of their actions (Fig. S1 and Tables S1 and S2) are complemented in ASMs by the exhaustive analysis of the modulation ranges caused by the allosteric mutations of all residues in the protein. As a result, the global picture of allosteric modulation, and, at the same time, all relevant details can be quantified and analyzed. For example, in addition to the dependence of the effect of mutation on a certain site averaged over corresponding sites in all domains/monomers (Fig.s S2-5), ASMs (Fig. 6) show how individual mutation affects the energetics of corresponding sites in every domain/monomer. The difference between regulation inside homological domains and subunits of oligomers also becomes visible. Additionally, inspection of ASMs allows one to determine mutations of which positions located outside regulatory exosite originate allosteric signaling similar to that caused by mutations in the regulatory site of interest, allowing thus, to obtain a required allosteric regulation caused by an alternative residue(s).

## Discussion

An existing consensus on the omnipresence and importance of allosteric mechanisms in regulation of functional activity of proteins and molecular machines (3, 11) and an increasing demand on the design of allosteric drugs (6, 7, 13, 16, 17) require an accurate theoretical understanding of allostery and development of computational models allowing the high-resolution control (on the level of individual amino acids or atomic groups) of allosteric signaling and regulation. In this work, we combine protein harmonic modelling with statistical mechanical formalism in a perturbation-based approach, which allows one to estimate per-residue allosteric free energy as a result of ligand(s) binding, mutation(s), and their combinations. We introduced the notion of allosteric modulation in order to distinguish targeted allosteric signaling to the sites/residues from the background effect caused by the same perturbation. We also formulated a notion of allosteric mutation, which we propose to consider as the basic element of protein activity modulation. We generically define the allosteric effect of mutation, modulation range, as the difference between the responses in regulated sites/residues obtained from stabilizing and destabilizing mutations, respectively. Finally, we also introduce the Allosteric Signaling Map (ASM) as an exhaustive description of allosteric signaling in the protein at per-residue resolution.

The generality of the model is shown here on three proteins with different degrees of oligomerization, functions, and mechanisms of regulation. Specifically, using example of homotetrameric PFK we analyze a switch between the activating and inhibiting modes originated by the binding of activator/inhibitor ADPa/ PEP in overlapping binding sites (27), as well as positive/negative cooperativity of their actions. The synergy of these effectors with a co-substrate ADPf and opposite modulation provided by Val54 and Arg154 mutations are also discussed. The ring-shaped tetrameric PGDH was used to show how domain-based allosteric regulation works and how mutations in hinges between regulatory and functional domains can affect the protein activity. The dependence of allosteric regulation on the location of mutations, such as mutation Ile365 and Leu370, and type of the amino acid substitution, *e.g*. stabilizing mutations Gly319/H2, Gly336/H4, and Gly337/H4, is also considered. IDE is a single-chain four-domain protein with the multi-component catalytic site allosterically regulated via binding to exosite (EXO) and ATP-binding (ATP) sites. A specific pattern of allosteric signaling with positive modulation in the diagonal domain and mixed – in flanking ones caused by sequential mutations in IDE domains was detected. We assume that this pattern of signaling can be explained by the structural homology of IDE’s four domains, whereas the diversity of sign and value of signaling in individual domains originates from the low sequence identity (at maximum 15 per cent) between them.

Examples of allosteric signaling considered here show that the diversity in modulatory modes of regulatory sites and competitiveness/cooperativity in their work can be allosterically affected by mutations located anywhere in the protein. Therefore, we introduced a description of the allosteric effect of mutation, modulation range, which is the difference between the effects of the stabilizing and destabilizing mutations on all residues of the protein (or its chain/subunit/domain). We argue that complete mutational scanning can be instrumental in taking the protein activity under allosteric control. The resulting Allosteric Signaling Map (ASM) can help to reveal a specific role of protein structural units (chains/domains) and associated critical length at which the mode switch in allosteric signaling takes place. It identifies strongest modulations, both positive and negative, inside these structures and between them. Finally, it allows one to infer alternative ways of allosteric regulation. Starting from the analysis of ASMs and comparison of effects of known allosteric sites and mutations, one can design new sites and use amino acid substitutions in order to obtain required mode and value of allosteric signaling to sites/residues of interest.

## Methods

Starting from the problems of causality and energetics in allosteric communication (24), we present a computational model with the goal to achieve a comprehensive allosteric control over protein activity.

Given a protein reference unbound/wild-type conformational state, denoted as “0” (unperturbed state), the C_α_ harmonic model energy function is

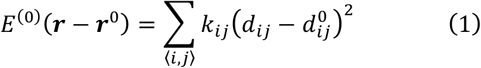

where ***r*** is the 3N-dimensional vector of coordinates of the C_α_ atoms, ***r***^0^ is the vector of C_α_ positions of the reference structure, *d_ij_* is the distance between the C_α_ atoms *i* and *j*, 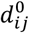 is the corresponding distance in the reference structure, and 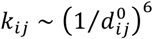 is a distance-dependent force constant with a global cutoff *d_c_* = 25 Å, with summation running over the pairs of neighbors 〈*i, j*〉 within the global distance cutoff (39, 40).

Let us now consider the protein in a perturbed state, which can be a combination of ligand binding and mutation events, respectively. The energy function associated with this perturbed *P* state is

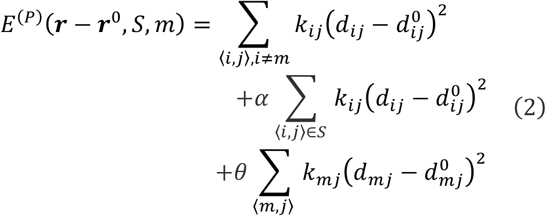

First, given the binding site *S* the protein ligand bound state is modeled by introducing an additional harmonic restraining term between all the residue pairs that compose the occupied binding site, with α is a stiffening parameter in Equation 4 (α=100, see (24) for details). Second, the protein mutated state is modeled by altering the strength of the force constants associated with the contacts between the mutated residue *m* and its neighbors. Two types of mutations are defined: UP (↑, stabilizing) mutation mimics a residue substitution with bulkier amino acids, a DOWN (↓, destabilizing) mutation - substitutions to small Ala/Gly-like residues. In Equation 2 the strength of the interactions between the residue of interest and its neighbors is scaled up by the factor θ =100 in case of UP mutation, while suppressed by a factor θ = 10^−2^ in case of DOWN mutations. Equation 2 considers for simplicity the case of single binding site and single-residue mutation and can be easily generalized to the case of multiple sites and mutations.

For the unbound/wild-type (0, Equation 1) protein state and the generally perturbed state (*P*, Equation 2), the Hessian matrix **K** = *∂*^2^*E* /*∂**r**_i_∂**r**_i_∂**r**_j_* is considered and the set of orthonormal normal modes 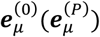, characterizing the configurational ensembles of two corresponding protein states are calculated. The modes ***e**_μ_* are used in the allosteric potential for evaluating the elastic work that is exerted on a particular residue *i* as a result of the ensembles described by the corresponding normal modes

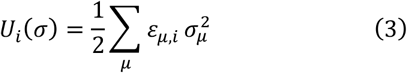

where *ε_μ,i_* are parameters defined from the normal modes as

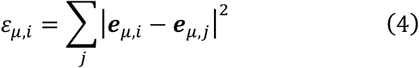

Essentially, the allosteric potential accounts for the change of displacement of the neighbors of residue *i* over the modes with a given set of Gaussian distributed amplitudes *σ* = (*σ*_1_, …,*σ_μ_*,…) with variance 1/*ε_μ,i_*. Since the generic displacement of a residue *i* can be written as 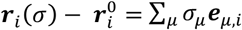, the vector *σ* represents a configurational state of residue *i*

Integrating over the ensemble of all possible configurations *σ* of a residue, the per-residue partition function is obtained

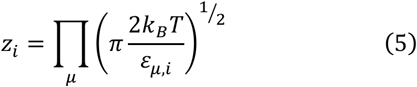

and, correspondingly, the free energy *g_i_* = −*k_B_* ln *z_i_* + *const*. Given the protein in a state *P* (bound, mutated, or their combination) the free energy difference with respect to the reference state 0 is

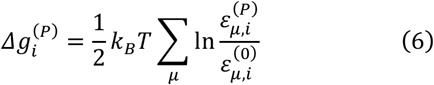

A positive value 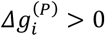 indicates that work exerted on residues *i* produces conformational changes as a result of a change in the ensemble of neighboring residues caused by the perturbation. A negative 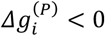 value shows a stabilization of residue *i*, which, on the contrary, may prevent it from the conformational change.

Cooperativity in the effect of perturbations in the case of oligomeric and multidomain proteins should also be considered. For instance, in case of a homo-oligomer with identical effector binding sites in each chain, one can define several bound states with an intermediate number of binding sites occupied (partially perturbed state *P_part_*) and one with all sites fully occupied (fully perturbed state *P_full_*). Thus, the cooperativity associated with the sequential perturbation of additional ligand binding sites can be obtained via the relation

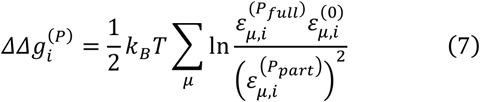

Equation 7 applies also in case of mutations.

In order to estimate a clean allosteric effect on a residue *i* due to the perturbation *P*, the deviation of the free energy difference 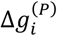 from its mean value over all the residues of the protein chain containing the residue *i* is considered

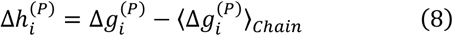

This background free energy change is called *allosteric modulation* 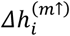 (positive or negative) caused by the perturbation *P*. The allosteric modulation, 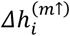 and 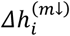, respectively, originated from UP and DOWN mutations can be generically calculated for any residue position in the protein. The difference

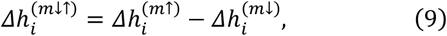

evaluates the allosteric *modulation range* caused by mutations from the smallest (Ala/Gly-like) to bulkiest residues. The allosteric modulation from the opposite mutation, from the largest to Ala/Gly-like residue, is, thus, 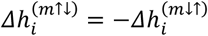. Finally, the allosteric modulation at the level of sites can be evaluated by averaging the per-residue modulations 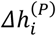 over the residues belonging to the site of interest

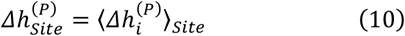

In order to compare calculated allosteric effects of mutations with experimental data and to put these data in the context of direct mutational effects, we considered PDZ domain as a widely studied protein (41) with well-documented allosteric regulation. Supp. Fig. S7 shows a comparison of direct and allosteric effects of mutations on the PDZ-domain’s binding site obtained with the model compared to the high-throughput experimental data on the effect of mutations on PDZ domain binding affinity.

